# Changes in structural network topology correlate with severity of hallucinatory behaviour in Parkinson’s disease

**DOI:** 10.1101/383141

**Authors:** Julie M. Hall, Claire O’Callaghan, Alana. J. Muller, Kaylena A. Ehgoetz Martens, Joseph R. Phillips, Ahmed A. Moustafa, Simon J. G. Lewis, James M. Shine

## Abstract

An inefficient integration between bottom-up visual input and higher-order visual processing regions is implicated in the manifestation of visual hallucinations (VH) in Parkinson’s disease (PD). Using graph theory, the current study aimed to investigate white matter contributions to this perceptual imbalance hypothesis. Twenty-nine PD patients reported their hallucinatory behaviour on a questionnaire and performed a behavioural test that has been shown to elicit misperceptions. A composite score derived from these measures was used as a proxy for hallucinations severity and was correlated to connectivity strength of the network using the Network Based Statistic approach. The results showed that the severity of VH was associated with reduced connectivity within a large sub-network. This network included the majority of the diverse club and showed overall greater *between*- and *within-module* scores, compared to nodes not associated with hallucination severity. Furthermore, a reduction in *between-module* connectivity in the lateral occipital cortex, insula and pars orbitalis, as well as decreased within-module connectivity in the prefrontal, somatosensory and primary visual cortices were associated with VH severity. In contrast, the severity of VH was associated with an increase in *between*- and *within-module* connectivity in the orbitofrontal and temporal cortex, as well as regions comprising the dorsal attentional and DMN. These results suggest that the severity of VHs is associated with marked alterations in structural network topology, highlighted by a reduction in connectivity strength across a large sub-network, as well as changes in participation across top-down visual processing centres, visual and attentional networks. Therefore, impaired integration across the perceptual hierarchy may result in the inefficient transfer of information that gives rise to VHs in PD.

## Introduction

Visual hallucinations (VHs) in Parkinson’s disease (PD) exist on a spectrum ranging from simple misperceptions, to complex well-formed images (1). With disease progression and loss of insight, VHs constitute a major source of distress for the patient (2, 3) and comprise a high degree burden for caregivers (4). Risk factors of VHs include older age and disease duration, sleep and mood disturbances as well as cognitive decline (5–7). Furthermore, previous work has shown that patients with VHs show disruptions in attentional processing (8), reduced performance on visuoperceptive tasks (9–11), decreased visual contrast sensitivity, colour discrimination (12) and acuity (13). Current models of VHs have therefore focused on the interaction of perceptual and attentional dysfunction (for review, see (14)). Specifically, it has been proposed that failure to effectively integrate information from different processing sites across the perceptual hierarchy is likely to contribute to VHs and misperceptions in PD (14–17).

Attention, prior experience and expectations strongly influence perception. Perceptual predictions, generated from a myriad of modalities across the brain, guide perceptual processes to facilitate the interpretation of noisy and ambiguous input (18–20). The orbitofrontal cortex (OFC) process coarse information projected from the visual cortex and provides an “initial guess” of an object’s identity (21). Previous work in PD patients with VHs has shown that the accumulation of sensory evidence is slow and inefficient, which may result in an over-reliance on these top-down predictions (22). Importantly, top-down visual processing regions can modulate neural activity in early visual regions, with expected stimuli leading to reduced activity (23). Additionally, activity within the default mode network (DMN), a network involved in mediating endogenous perception, has shown to be increased during a misperception in this patient population (24). Therefore, VHs may arise when perceptual input is not properly integrated and internally generated images interfere with the perceptual process (22, 25–27).

While functional neuroimaging studies have made significant contributions to our understanding (24, 28–31), less is known about the involvement of white matter changes in the manifestation of VHs in PD. Experiments using diffusion tensor imaging (DTI) have reported altered white matter integrity in the optic nerve and optic radiation (32) as well as ascending tracts from the cholinergic nucleus basalis of Meynert to parietal and occipital cortical regions (33). However, given the involvement of large-scale brain networks in perception, unique insights into white matter changes associated with VHs can be gained by investigating whole brain network topology. Topological features of the human connectome allow us to describe the arrangement of connections within and between segregated sub-modules (34). Specifically, nodes that integrate these specialist communities are crucial for incorporating information streams of different modalities, which is essential for processes such as perception (35, 36). Therefore, investigating network topology can provide novel insights in changes across different perceptual hierarchies.

The current study aimed to examine whether VHs are associated with changes in structural network topology. We hypothesized that the severity of hallucinatory behaviour would be associated with ineffective information processing as shown by reduced *between-module* scores in visual networks, reflecting reduced visual input to integration centres. Furthermore, increased *between-module* scores across top-down perceptual prediction areas and the DMN could indicate an over-reliance on regions involved in the generation of internal percepts (37).

## Methods

Twenty-nine patients with idiopathic PD were included in this study. Demographic information including age, disease duration and levodopa dose equivalent (LEDD) were obtained for all participants. All patients were assessed on the Hoehn & Yahr clinical stage (38) and the motor aspect of the MDS-UPDRS (part III) (39). Global cognition was assessed using the Mini-Mental State Examination (MMSE) (40) and set-shifting performance was assessed using the Trail Making Test part B minus part A (TMT_B-A_)(41). The study was approved by the local ethics committee and was in accordance with the principles of the Helsinki Declaration. Written informed consent was obtained from all participants before participation.

### Bistable Percept Paradigm

All patients performed the Bistable Percept Paradigm (BPP) (17), a behavioural task capable of inducing misperceptions in susceptible patients. In this task, patients were presented with either single or bistable percepts (i.e. “hidden” images as shown in Figure 1) for a maximum of 30 sec in a randomised order. The participant had to decide whether the stimulus was a single or hidden image by a button press and describe to the examiner what they had seen. The recorded responses included the following: 1) correct single or correct hidden, 2) “missed”, recorded when the subject perceived a single image when a bistable percept was presented and 3) “misperceptions”, recorded when a subject incorrectly identified a single image as a bistable image, i.e. incorrectly reported an image that was not presented on the screen.

**Figure 1.**
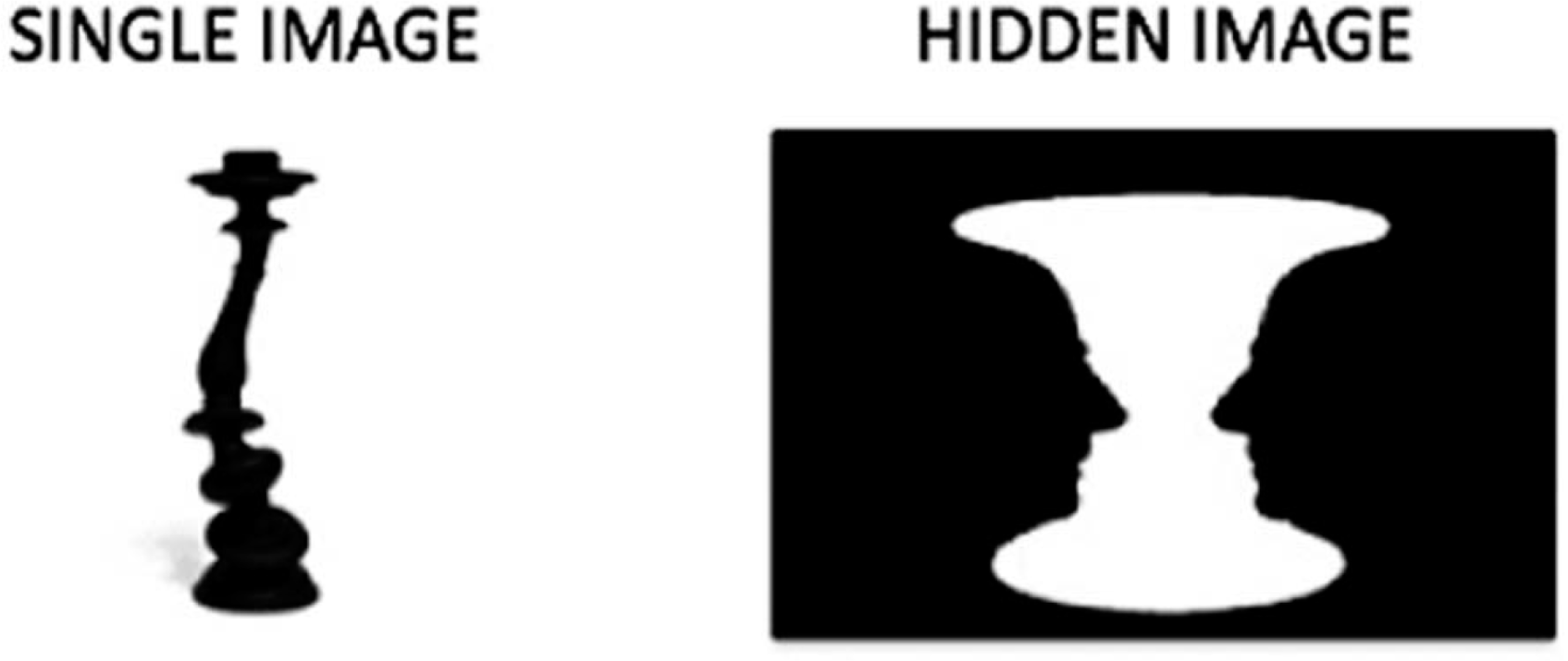
Example of single and hidden images of the BPP (42)

### Psychosis and Hallucinations Questionnaire

All patients completed the Psychosis and Hallucinations Questionnaire (PsycH-Q) (43). This validated questionnaire consists of two parts, of which the responses of the first part (PsycH-Q_A_) were included in this study. The PsycH-Q_A_ consists of three subscales including 1) visual misperceptions, which includes questions about the presence of VHs, passage hallucinations and three frequently reported contents of VHs including people, animals and objects; 2) sensory misperceptions, including audition, touch, olfaction, and gustation, and; 3) disordered thought and psychotic behaviour. Participants rated the frequency of their symptoms on a 5-point Likert scale, ranging from 0 (“never experienced”) to 4 (“experienced daily”). The total score was calculated by summing the responses (43) (see Supplementary Materials). Part B of the PsycH-Q assesses symptoms related to VHs (i.e. attention and sleep) and was not included in this study.

### Composite score

The percentage of misperceptions on the BPP and the total score on the PsycH-Q_A_ were standardized and then summed to create a composite score that reflected the severity of visual hallucinatory behaviour (hereafter referred to as the hallucination severity score (HSS)). The HSS was correlated with the demographic variables using parametric or non-parametric correlations depending on the distribution of the variables and was used as a correlate in the imaging analysis.

### MRI Acquisition

All participants underwent magnetic resonance imaging (MRI) using a 3-Tesla General Electric Discovery MR750 scanner (GE Medical Systems) with an 8-channel phased array head coil. Diffusion weighted images (DWI) were obtained by using echo-planar imaging sequences with 61 different motion probing gradient directions (TR/TE: 7025/80 ms, 55 transverse slices, slice thickness: 2.5 mm, matrix: 256 × 256, FOV: 240 x 240 mm). The effective diffusion weighting was b = 1000 s/mm^2^ and four volumes with no diffusion weighting (b = 0 s/mm^2^) were obtained at the beginning of each diffusion sequence. 3D T1-weighted, anatomical images were obtained (TR/TE/TI: 7.2/2.7/450 ms, voxel size 1 × 1 × 1 mm, 196 transverse slices, 256 × 256 matrix, FOV: 256 x 256 mm, flip angle 12°). The 3D-T1 images were used for individual registration between T1 weighted anatomical and the DWI images and cortical parcellation using FreeSurfer (version 5.3; http://surfer.nmr.mgh.harvard.edu).

### Diffusion tensor imaging pre-processing and deterministic fibre tracking

DTI pre-processing was performed using the FMRIB Software Library (FSL, http://fsl.fmrib.ox.ac.uk). The pre-processing steps were as follows: *i)* DTI images were corrected for susceptibility, head motion and eddy current induced geometrical distortions using FSL’s tool *eddy*; *ii)* a binary brain mask was created using *bet; iii)* images were realigned using a rigid body registration to the b = 0 image; then *iv)* a tensor was fitted in each voxel (44); followed by *v)* the computation of the fractional anisotropy (FA) level based on the eigenvalues for each voxel, in order to determine the preferred diffusion direction within a voxel. FA thus serves as a surrogate measure of white matter integrity, with lower levels of FA reflecting reduced white matter integrity (45–47). The preferred diffusion direction information was then used to reconstruct the white matter tracts of the brain using a deterministic tracking approach based on the *fibre assignment by continuous tracking* (FACT) algorithm (48). Deterministic tractography yields less false positive tracts compared to probabilistic methods (49). False positives are detrimental in network modularity as they occur more prevalently between than within modules (50). A streamline was started from eight seeds within each voxel of the brain (grey and white matter) following the main diffusion direction of the voxel and stopped when: *i)* the FA value < 0.1; *ii)* the traced fibre made a turn >45°; or *iii)* when the tract left the brain mask. The images were acquired when reverse phase-encoding direction approaches were not the standard procedure within acquisition protocols, which could have influenced the registration of diffusion and anatomical images. Therefore, anatomically constrained tractography was not applied (51). The weighted brain network was calculated for each participant and consistency thresholding at 50% was applied (i.e. including the tracts found in 50% of the patients) (52). The mean density of the thresholded group matrix was 8.7%. To verify the results were not skewed by the choice of threshold, we also applied the thresholding method that retained most consistent edges across subjects but controlling for their distance (i.e. the consistency of edges within “bins” based on their length to avoid preferential retention of short edges (53)). The mean density of the group matrix using this threshold was 13.2%.

### Network Based Statistic

A Network Based Statistic (NBS) analysis was applied to investigate whether the HSS was associated with altered connectivity strength in an interconnected subnetwork of the brain (54). NBS is a non-parametric method for connectome-wide analysis, which aims to detect specific pairs of brain regions showing a significant effect of interest, while controlling for family-wise-error (FWE) rate. Importantly, no inferences of individual connections are made; instead the null-hypothesis can only be rejected at the sub-network level. As such, NBS is similar to the cluster-based multiple-comparisons approaches used in standard functional MRI analysis. To identify changes in sub-networks associated with the HSS, the t-statistic was set at 1.7 determined using the critical value of the t-distribution for our sample size (55). Connections were deemed significant at FWE-corrected *p*-value < 0.05 (one-sided) using 5000 permutations.

To investigate whether the sub-network involved particular functional networks, we investigated whether nodes in the sub-network that correlated with the HSS overlapped with previously defined resting state networks. To this end, seven canonical resting state networks from the Yeo *et al*. atlas (56) were overlaid with the structural parcellation and the percentage of nodes from each network included within the structural sub-network that inversely related to HSS was calculated for each resting state network. To analyse whether this overlap occurred significantly above chance, we randomly permuted the resting state network identity of each region (5000 iterations) and used the overlap between the randomized vector and the original node assignment to populate a null distribution. To test whether each individual resting state network overlapped with the significant sub-network, their overlap was compared with the null distributions. A resting state network was identified as targeted if the true overlap was more than the 97.5^th^ percentile of null distribution (i.e. the top 2.5%). A network was considered not to be associated with the HSS if the overlap was less than the 2.5^th^ percentile of the null distribution.

### Graph theoretical analysis

The graph organizational measures were computed using the Brain Connectivity Toolbox (http://www.brain-connectivity-toolbox.net) (57). The thresholded, weighted brain networks were then partitioned into modules, which are non-overlapping groups of highly connected nodes that are only sparsely connected with other modules, using the Louvain algorithm (57). To account for the stochastic nature of the Louvain algorithm, a consensus partition was identified by calculating the module assignment for each node 500 times. To define an appropriate value for the resolution parameter (γ), the Louvain algorithm was iterated 100 times across a range of values (0.5 – 2.0 in steps of 0.1) of the group mean connectivity matrix and then estimated the similarity of the resultant partitions using mutual information. The γ parameter of 1.9 provided the most robust estimates of topology across the iterations and was used to determine the optimal resolution of the network modularity.

After the nodes were assigned to their modules, their intra- and inter-modular connectivity were calculated. Intra-modular connectivity was calculated using the module degree z-score *W_i_* (see equation 1), in which a positive score reflects high *within-module* connections (compared to the node’s average number of connections), and negative z-scores denote the opposite. Inter-modular connectivity was calculated using the participation coefficient *B_i_* (see equation 2). Low *B_i_* values indicate few *between-module* connections, whereas high *B_i_* values indicate uniformly distributed connections across modules (github.com/juliemaehall/topology). High *W_i_* and high *B_i_* scores are not mutually exclusive (58).

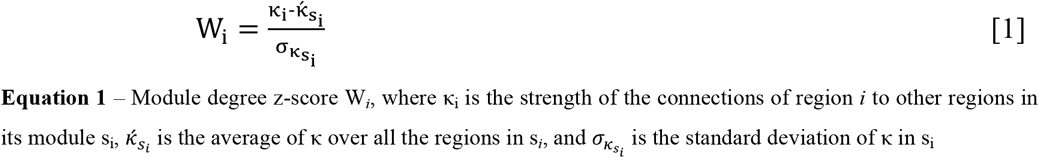

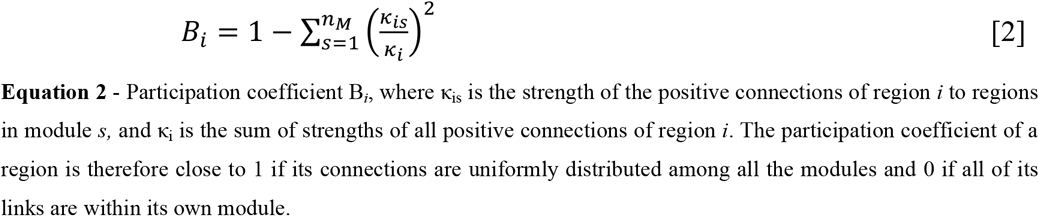

To test whether nodes within the sub-network identified using the NBS analysis differed from nodes not included in the sub-network, the average *W_i_* and *B_i_* of the subnetwork were contrasted against the average *W_i_* and *B_i_* of the nodes not included in the sub-network using non-parametric permutation testing.

To test whether the HSS correlated with the *W_i_* and *B_i_* nodes across the whole brain connectome, a Spearman’s rho correlation was performed followed by a non-linear permutation test using 5000 iterations to control for multiple comparisons (59), using an alpha of 0.05. This approach was repeated using the different threshold (53) and the outcome was correlated to the *W_i_* and *B_i_* using the original threshold. Both the *W_i_* and *B_i_* scores calculated using the aforementioned threshold highly correlated with the *W_i_* and *B_i_* scores calculated with the consensus threshold (*r* = 0.92 *and r* = 0.94, respectively), indicating that the results were not biased by the chosen thresholding method.

### Diverse club analysis

We identified the ‘diverse club’ of the network, which comprised of the top 20% of *B_i_* nodes (60). These nodes play an important role in network integration and changes to these nodes could affect between module communication (60). We normalized the diverse club coefficient in reference to a null model: a random vector with a preserved modular structure was created by randomizing the mean participation coefficient of each node for 5000 iterations. The diverse club was identified as those regions with a participation coefficient greater than the 95^th^ percentile of the permuted distribution. We investigated whether the number of diverse club nodes was significantly higher within the sub-network associated with the HSS, compared to nodes that were not included in the sub-network identified using the NBS analysis.

## Results

### Demographics

Table 1 presents the descriptive variables of the 29 patients. The mean percentage of misperceptions on the BPP was 18.48 (range: 0 – 49), and the mean score on the PsycH-Q_A_ was 9.48 (range: 0 – 34 (max score = 52)), highlighting a diverse range of hallucinatory behaviour in the patient cohort. Finally, to verify the severity score to the ‘gold standard’, we correlated the HSS in a large cohort of patients with PD and Lewy Body Dementia (n = 75) to the MDS-UPDRS item 2 and found a correlation of *r* = 0.53 (*p* < 0.001). However, given higher construct validity (43), we opted to use the PsycH-Q_A_ scores for the remainder of our analysis.

**Table 1.** Demographics and clinical variables

| Variable | Mean (range) | Correlation with HSS |
| --- | --- | --- |
|  |  | r (p-value) |
| Age (y) | 65.1 (51 – 84) | 0.18 (0.345) |
| Duration (y) | 5.8 (1.2 – 16) | -0.05 (0.785) |
| LEDD | 656.9 (125 – 1548) | -0.06 (0.767) |
| H&Y | 1.9 (1 – 3) | 0.01 (0.949) |
| MDS-UPDRS III | 25.5 (7 – 55) | 0.29 (0.125) |
| MMSE | 28.5 (25 – 30) | -0.06 (0.762) |
| TMT<sub>B-A</sub> | 78.9 (-1 – 123) | 0.33 (0.08) |
| BPP % misperceptions | 18.48 (0 – 49) | - |
| PsychH-Q<sub>A</sub> | 9.48 (0 – 34) | - |

LEDD = Levodopa equivalence daily dose; MDS-UPDRS III = motor part of the Movement Disorder Society Unified Parkinson’s Disease Rating Scale; H & Y = Hoehn and Yahr; MMSE = Mini Mental State Examination; TMTB-A = Trail Making Test part B – part A; BPP = Bistable Percept Paradigm; PsychH-Q<sub>A</sub> = Psychosis and Hallucinations Questionnaire, part A

The HSS showed a positive correlation trending towards significance with the TMT_B-A_ (*r* = 0.33, *p* = 0.08). No significant correlations were observed between the HSS and other demographic and clinical variables.

### The HSS correlated with decreased connectivity in a large sub-network

As illustrated in Figure 2, the NBS analysis revealed a sub-network comprising 183 edges (12% of the edges in the thresholded connectivity matrix) and 127 nodes with reduced FA-based connectivity strength correlated to the HSS (*p* < 0.05). The effects presented with a fairly liberal threshold, suggesting the changes related to the HSS are subtle yet topological extended (54). No significant sub-network was identified in the opposite direction (positive correlation between the HSS and connectivity strength).

**Figure 2.**
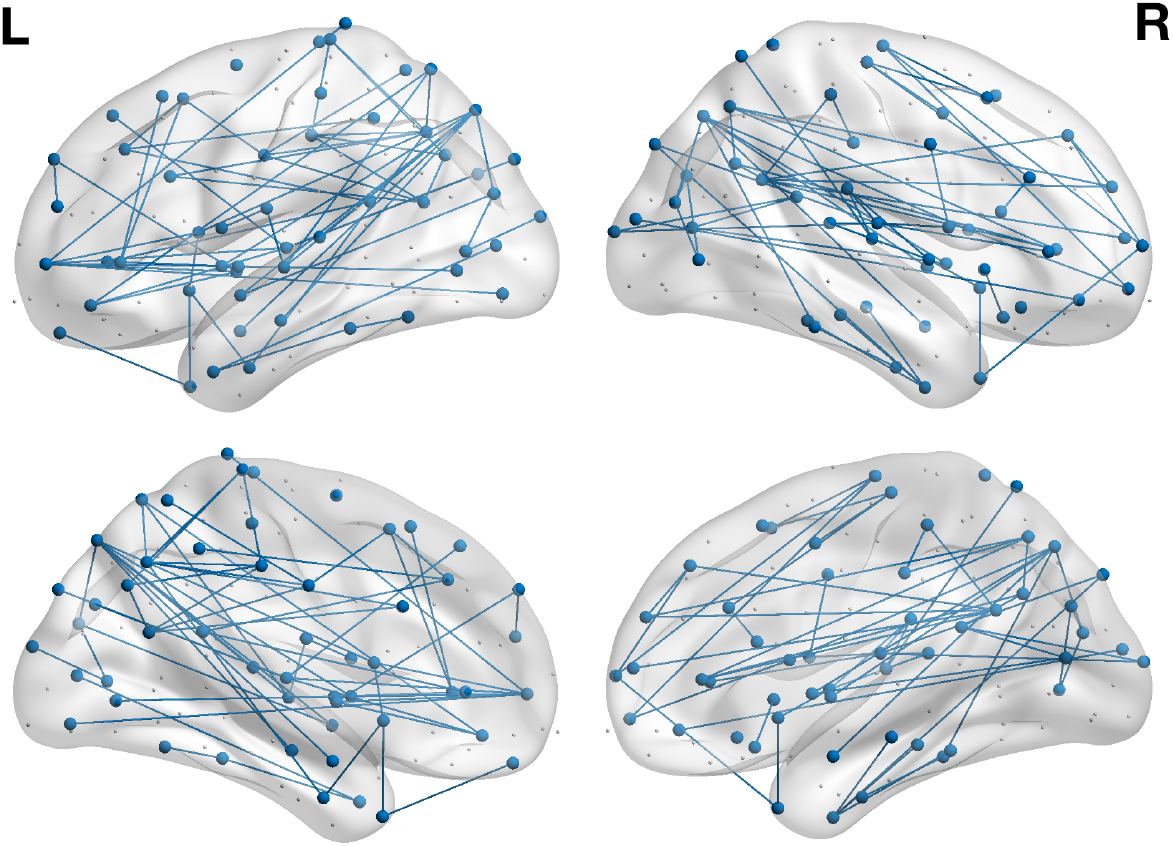
NBS analysis reveals a sub-network, comprising 183 edges and 127 nodes with reduced connectivity strength correlated to increased HSS (p < 0.05). Figure visualized with BrainNet Viewer (61).

Furthermore, the average *W_i_* and *B_i_* scores within the sub-network were 0.149 and 0.759 respectively. These values were significantly higher (*p* < 0.05) than the average *W_i_* and *B_i_* scores of the nodes outside the network (−0.178 and 0.628, respectively).

### The sub-network includes all sub-cortical nodes but did not target a specific cortical resting state network

The sub-network that showed decreased connectivity strength correlated with the HSS included the majority of the sub-cortical nodes (*p* < 0.05). As illustrated in Figure 3, the sub-network further included nodes across the cortex, however the somatomotor network was relatively spared (*p* < 0.05). Sixty-five percent of the nodes in the DMN were part of this sub-network, however this was not significantly above chance (*p* > 0.05).

**Figure 3:**
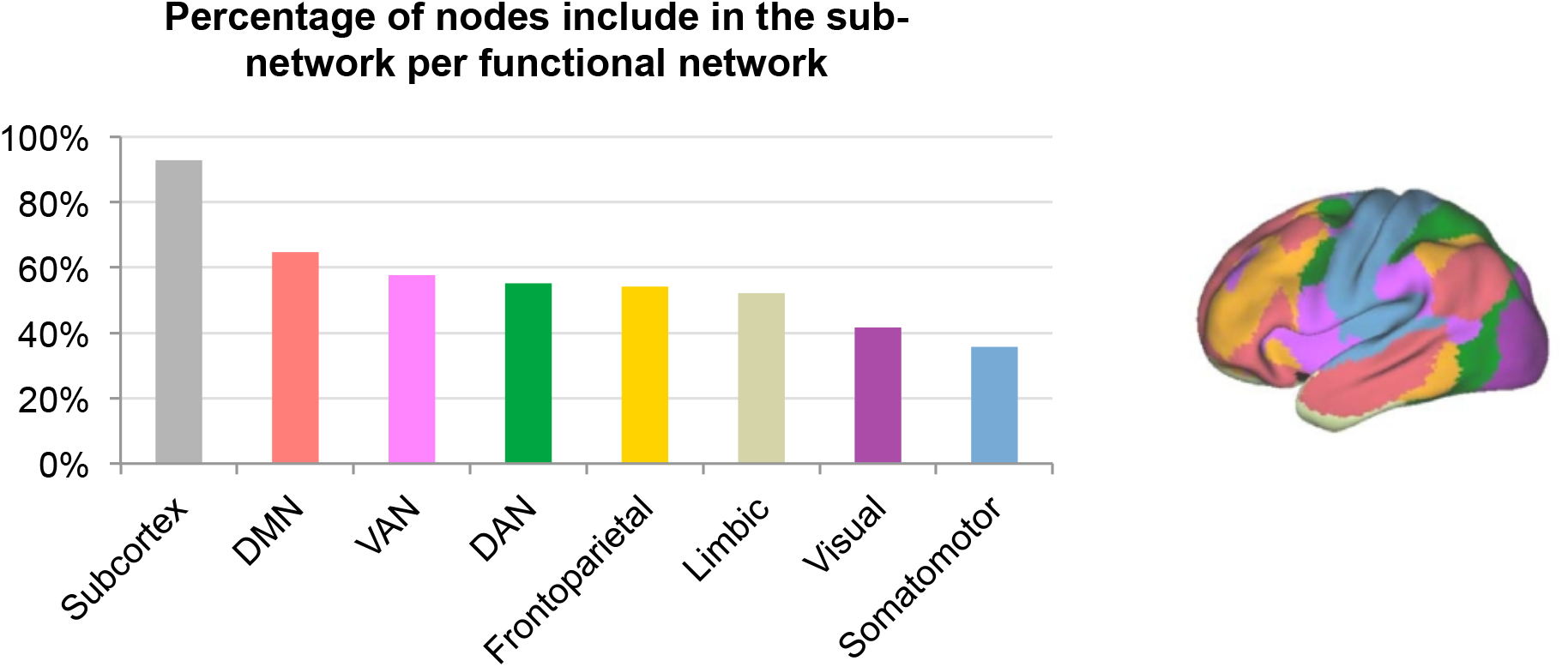
Left panel: the percentage of nodes included in the sub-network for each resting state network. Right panel: the functional resting state networks of the Yeo atlas (56). DMN = default mode network; VAN = ventral attentional network; DAN = dorsal attentional network.

### Nodes in the sub-network show high participation scores

Eighteen nodes were included in the diverse club (see Supplementary Materials). Seventeen of the eighteen nodes (94%) of the diverse club were included in the aforementioned sub-network, which was deemed significantly above chance (p < 0.001). As illustrated in Figure 4, nodes with high participation coefficients were more often part of the sub-network.

**Figure 4.**
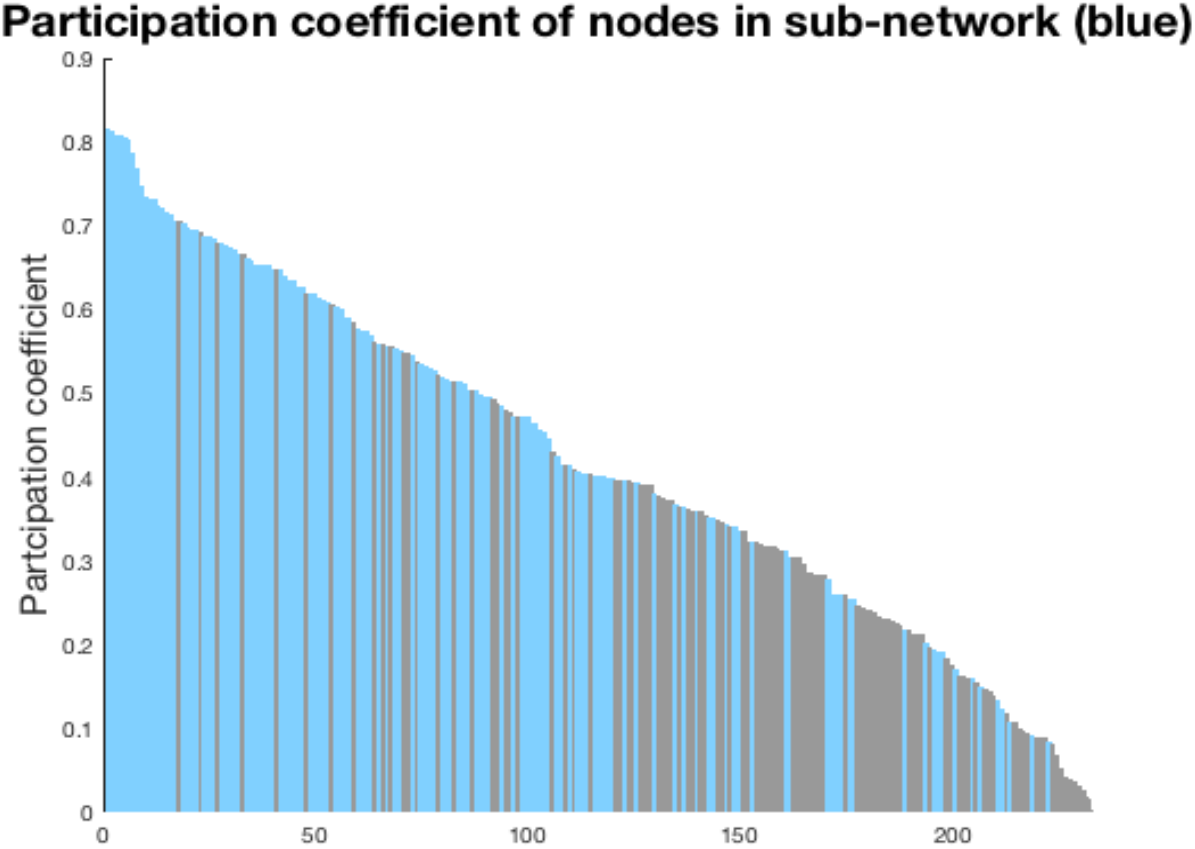
Nodes ranked according to the B_i_ scores. Blue: nodes included in the sub-network; grey: nodes not included in the sub-network correlated to the HSS.

### The HSS correlated with W_i_ and B_i_ scores

When investigating the whole structural connectome, the HSS positively correlated to regional *B_i_* (i.e. higher participation scores were associated with higher severity values) for nodes in the left medial OFC, a node in the right anterior and left posterior cingulate, precuneus, and the caudal middle frontal gyrus. Furthermore, nodes in the right occipital, pars orbitalis and insula showed negative correlations between the HSS and participation coefficient (i.e. lower participation scores were associated with higher scores on the HSS; see Table 2 and Figure 5).

**Table 2.**
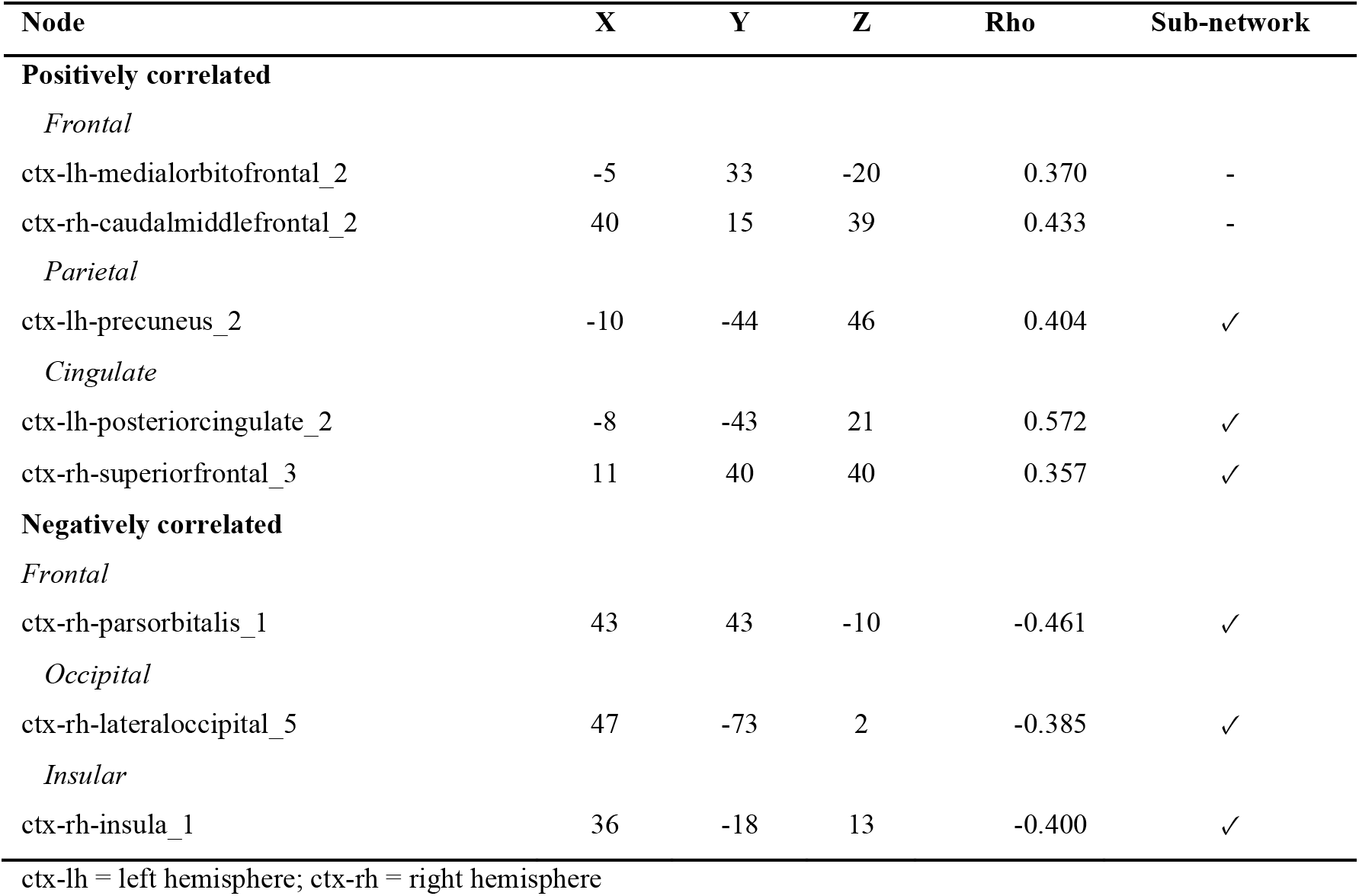
Spearman’s rho correlation between the participation coefficient and the HSS (*p* < 0.05; permutation test)

**Figure 5.**
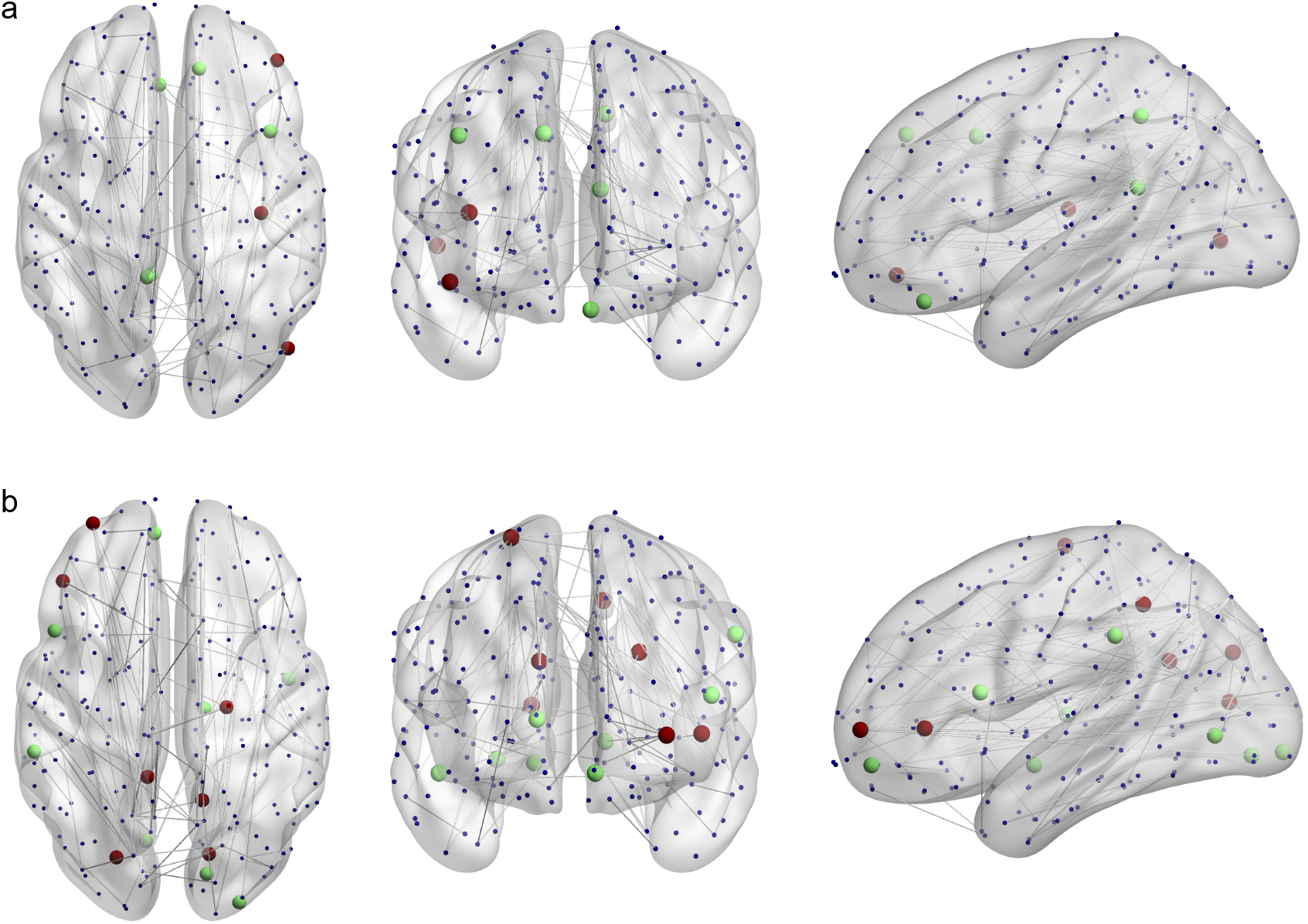
a) Nodes with a significant correlation between participation coefficients and the HSS; b) Nodes with a significant correlation between the module degree z-score and HSS. Green indicates a positive correlation, red indicates a negative correlation.). Figure visualized with BrainNet Viewer (61).

Increased HSS scores were further associated with increased module degree z-scores in the right thalamus, the bilateral lingual, left medial OFC, pars opercularis and supramarginal gyrus, the right lateral occipital and superior temporal cortices. Decreased HSS were associated with increased module degree z-scores in the bilateral precuneus, the left parts triangularis, rostral middle frontal and superior parietal cortex, the right pericalcarine and precentral gyrus (see Table 3 and Figure 5).

**Table 3.**
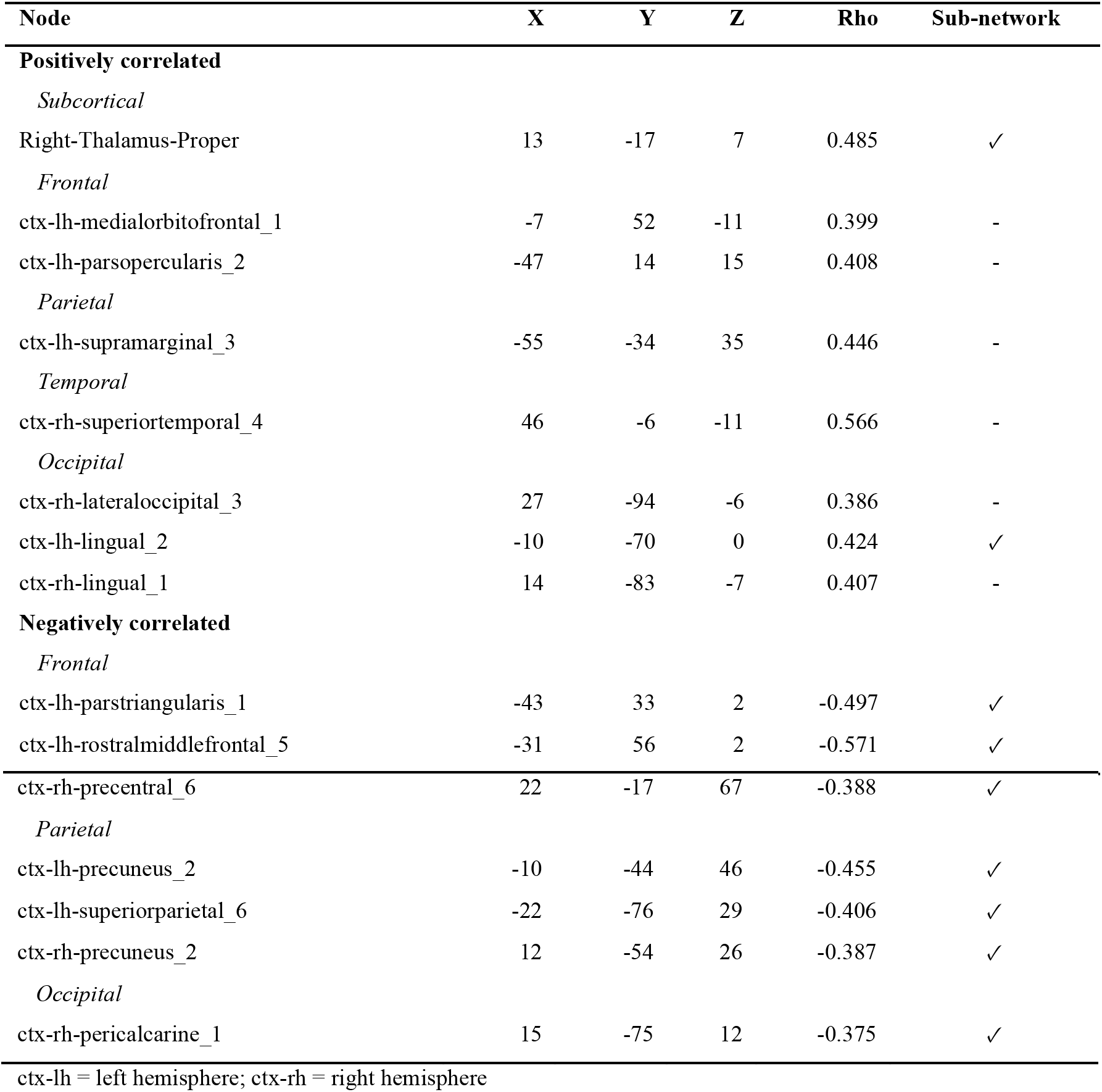
Spearman’s rho correlation between the module degree z-score and the HSS (*p* < 0.05; permutation test)

## Discussion

The aim of this study was to determine whether changes in structural network topology were associated with hallucinatory behaviour in PD. We showed that severity of hallucinatory behaviour was correlated with reduced connectivity across a bilateral sub-network. Regions within this sub-network showed higher participation and module degree z-scores compared to regions outside this network. The loss of connectivity strength may force the system to adapt and reroute information across less efficient pathways, impeding the standard sensory integration process. Importantly, 94% of the nodes in the diverse club were included in this sub-network. This community of high participation nodes is thought to control the integration of relatively segregated regions (60). Indeed, the diverse connectivity pattern of these nodes makes them crucial for the functional coordination of brain regions during tasks, and activity in these nodes predicts changes in the coupling of other regions (60). Severity of hallucinatory behaviour may thus be the result of impaired integration and segregation of brain networks or ‘modules’, affecting effective information transfer. Finally, we showed regional changes in participation associated with hallucination severity (the HSS score), with a positive correlation between participation scores in the medial OFC, cingulate, precuneus and middle frontal gyrus and the HSS and negative correlation with participation scores in the lateral occipital cortex, pars orbitalis and insula. These findings suggest a reweighting of the regions along the perceptual hierarchy, which may give rise to VHs.

Lower participation of the lateral occipital cortex may reflect reduced early visual processing, resulting in ineffective accumulation of visual information from the environment. Previous work using a Bayesian drift diffusion model has demonstrated that accumulation speed and quality of perceptual information are reduced in PD patients with VH (22). Furthermore, reduced quality or integration of visual stimuli may increase perceptual uncertainty, a suggestion that aligns with increased participation in the dorsal anterior cingulate cortex (62). Perceptual uncertainty may place excessive emphasis on top-down prediction centres, which subsequently could lead to a reduced activity in early visual regions (23).

This emphasis on top-down visual processing centres is supported by the increased participation coefficient and module degree z-score in the medial OFC. The OFC has an integrative function across brain networks as evidenced by its high participation coefficient. Additionally, this region is thought to facilitates recognition during visual perception by integrating incoming sensory information with previous experiences and expectations (63). During typical visual perception, the OFC is activated early in response to visual stimuli, receiving low spatial frequency signals from the visual cortex (64). Notably, only stimuli resembling known objects are shown to activate the OFC, which in turn generates a semantic association and provides a predictive signal to the visual system (65). Conversely, visual stimuli that carry no meaningful association do not activate the OFC in healthy individuals. Hence, it could be speculated that due to decreased quality of visual input, inappropriate recruitment of the OFC occurs, which may result in ascribing false associative information to visual stimuli.

The manifestation of VHs has previously been recognized as a dysfunction between the attentional networks (31). Specifically, patients with VHs are shown to be less able to recruit the dorsal attentional network (DAN), which enables the selection of appropriate sensory stimuli (66). With reduced control of this network, ambiguous stimuli might instead be interpreted by the ventral attentional network (VAN) and the DMN, which are less well equipped for this task. Our results showed increased participation in the posterior cingulate cortex (PCC), a key hub of the DMN. PCC activity has been implicated in regulating the focus of attention, specifically, the shift from the external world into internal mentation (67). Furthermore, the PCC is involved in mind wandering and supports internally directed cognition (68). A failure to suppress PCC activity may lead to the intrusion of internal thoughts into task performance (69). Moreover, a positive correlation was found between the HSS and the module degree z-score of the left pars opercularis, a node in the VAN, a network that is activated when expectations in perception are violated (24, 70). Conversely, a negative correlation between the HSS scores and module degree score and participation coefficients was found in other nodes of the VAN, namely the left pars triangularis, the right pars orbitalis and insula. The left pars triangularis supports resolving competition between simultaneously active representations (71), whilst the insula plays an important role in dynamically shifting attention between the attentional control networks (72). The anterior insula has previously been shown to be reduced in volume in PD patients with VH (24, 73). Together, these results suggest that ineffective communication between attentional networks in the brain may predispose an individual to hallucinate. Surprisingly, the participation coefficient of a node within the DAN (‘ctx-rh-caudalmiddlefrontal_2’) showed a positive correlation with the HSS. This node was not part of the sub-network and it may be possible that this is a compensatory response to the loss of connectivity strength in the other DAN regions. Notably, the connectivity matrix shows between module connections of this region with nodes in the somatomotor and the frontoparietal network, but not with the DMN or VAN.

Finally, all nodes that showed negative correlations with the HSS were included in the sub-network. Decreased *within-module* scores were found across the prefrontal and the somatosensory association cortex, as well as in the primary visual cortex, whilst there was a positive correlation between the HSS and the bilateral secondary visual cortex, perhaps as a result of the decreased visual input from V1. Additionally, the supramarginal gyrus, a node that has been shown to be functionally active during spatial perception but also during visual imagery (74) showed an increased module degree z-score with increasing severity of VHs. Furthermore, a positive correlation with the HSS and the module degree score in the superior temporal cortex, a region involved in auditory processing was found. Although further research is needed, it could be speculated that increased visual uncertainty may stimulate other sensory processing areas.

This study has several limitations worth noting. Firstly, the DWI data was acquired without EPI distortion correction. This may have affected the accuracy of registration between DWI and T1 images in the frontal and temporal cortices. Due to relatively low diffusion weighting used in the current MRI protocol, it was chosen to employ DTI rather than more sophisticated methods such as constrained spherical deconvolution, an algorithm that more adequately deals with multiple fibre directions within one voxel than DTI. Furthermore, as the investigation was conducted in a relatively small group of PD patients, future studies should replicate our findings in a larger sample size.

## Conclusions

We conclude that hallucinatory behaviour in PD patients is associated with marked alterations in structural network topology. Severity of hallucinatory behaviour was associated with decreased connectivity in a large sub-network that included the majority of the diverse club. These changes may result in an inefficient rerouting of information across less efficient pathways, which may lead to impaired visual integration processes. Furthermore, nodes within the orbitofrontal cortex and temporal lobes showed increased participation scores, whilst the visual association cortex, insula and middle frontal gyrus showed decreased scores associated with the HSS score. These findings suggest that impaired integration across different regions along the perceptual hierarchy may result in inefficient transfer of information. A failure to effectively switch between attentional networks and the intrusion of internal percepts could give rise to perceptual glitches, such as misperceptions and hallucinations.

## Acknowledgements

We thank the patients and their families who contribute to our research at the Parkinson’s Disease Research Clinic. We thank Dr Váša for sharing his thresholding code (github.com/frantisekvasa/matlab_general). The DWI data was processed during the 10kin1day initiative at the Dutch Connectome Lab. This research was supported by Sydney Informatics Hub, funded by the University of Sydney. JMH is supported by a Western Sydney University Postgraduate Research Award; CO is supported by a National Health and Medical Research Council Neil Hamilton Fairley Fellowship (1091310) and by the Wellcome Trust (200181/Z/15/Z). AJM is supported by an Australian Postgraduate Award at the University of Sydney; KAEM is supported by a University of Sydney Postdoctoral Fellowship; JRP is supported by an Australian Postgraduate Award at the Western Sydney University. SJGL is supported by National Health and Medical Research Council-Australian Research Council Dementia Fellowship (#1110414) and this work was supported by funding to Forefront, a collaborative research group dedicated to the study of non-Alzheimer disease degenerative dementias, from the National Health and Medical Research Council of Australia program grant (#1037746 and #1095127). JMS is supported by a National Health and Medical Research Council CJ Martin Fellowship (1072403). AAM has no funding source to disclose.

